# Pan-cancer analysis reveals the prognostic potential of THAP9/THAP9-AS1 Sense-Antisense gene pair in human cancer

**DOI:** 10.1101/2022.01.20.477092

**Authors:** Richa Rashmi, Sharmistha Majumdar

## Abstract

Human THAP9 is a domesticated transposon homologous to the Drosophila P-element transposase (DmTNP). However, the exact functional role of THAP9 is unknown. THAP9-AS1 (THAP9-anti sense1) is a lncRNA involved in various cancers. Together they are arranged head-to-head on opposite DNA strands forming a sense and anti-sense gene pair. No studies have analyzed THAP9 and THAP9-AS1 from a pan-cancer perspective. First, we analyzed the bidirectional promoter region between THAP9 and THAP9-AS1 for core promoter elements and CPG Islands. Moreover, we analyzed the expression, prognosis, and biological function of the two genes across different types of tumors included in The Cancer Genome Atlas and GTEx datasets. The results showed that the expression of both genes varies in different tumors. Expression of the THAP9 and THAP9-AS1 gene pair was observed to be strongly correlated with prognosis in patients with tumors; higher expression of the two genes was usually linked to poor overall and disease-free survival. Therefore, they may serve as a potential biomarker of clinical tumor prognosis. Further, we performed gene coexpression analysis using WGCNA followed by differential gene correlation analysis (DGCA) across 22 cancers to study which genes share the expression pattern THAP9 and THAP9-AS1 and if their interacting partners change in various conditions. Gene ontology & KEGG pathway analysis of genes coexpressed with the two genes identified gene enrichment related to DNA binding and Herpes simplex virus 1 infection.

## Introduction

If the 5’ ends of the two genes are adjacent to one another on opposite DNA strands, and the two genes are transcribed divergently, they are called head-to-head genes. The region between the TSS of head-to-head genes can be identified as a putative bidirectional promoter. Eukaryotic genes are sometimes organized in a head-to-head architecture, sharing a bidirectional promoter region for regulating the expression of the two genes (Hurst et al., 2004). Genome-wide analysis has shown that more than 10% of the human genes are arranged in a bidirectional head-to-head architecture with their transcription start sites located <1 kb apart (Adachi and Lieber, 2002; Li et al., 2006; Trinklein et al., 2004).

The transcriptional regulation of genes that share a bidirectional promoter is complex. They can be positively correlated, such as human collagen genes COL4A1/COL4A2 (Burbelo et al., 1988; Heikkilä et al., 1993), negatively correlated such as mouse TO/KF (Schuettengruber et al., 2003), or they can show tissue-specific or condition-specific correlation such as human HSP60/HSP10 which are coordinated to respond to induction signals (Hansen et al., 2003). Interestingly, several recent genome-wide studies have reported that sense genes positively correlate with antisense genes in the same tissues or cells. For instance, it has been (Balbin et al., 2015) observed that 38% of annotated antisense RNA transcripts positively correlated with sense gene expression in 376 cancer samples comprising nine tissue types. Moreover, several bidirectional gene pairs are associated with human diseases, such as BRCA1/NBR2 (Auriol et al., 2005), ATM/NPAT (Luo et al., 1998), DHFR/MSH3 (Shinya and Shimada, 1994), and SERPINII/PDCD10 (Chen et al., 2007).

The THAP9 and THAP9-AS1 genes are a putative bidirectional gene pair; Their transcription start sites are located 166 bases apart, and they are arranged in a head-to-head or divergent manner on opposite DNA strands. THAP9 is a domesticated human DNA transposase, homologous to the widely studied Drosophila P-element transposase (Majumdar et al., 2013). The THAP9 protein shares 40% similarity to the P-element transposase and probably does not transpose *in vivo* due to the absence of terminal inverted repeats and target site duplications. Despite being domesticated, it has retained its catalytic activity (Majumdar et al., 2013; Majumdar and Rio, 2015). hTHAP9 belongs to the THAP (Thanatos-associated protein) protein family in humans, with twelve members (hTHAP0-hTHAP11). All human THAP proteins are characterized by an amino-terminal DNA-binding domain called the THAP domain, typically 80-90 amino acid residue-long, and possesses a C2CH Type Zinc Finger (Campagne et al., 2010; Sabogal et al., 2010).

Many THAP family proteins are known to be involved in human diseases. THAP1 has been associated with DYT6 dystonia (a hereditary movement disorder involving sustained involuntary muscle contractions) (Sengel et al., 2011). Regulation of THAP5 by Omi/HtrA2 has been linked to cell cycle control and apoptosis in cardiomyocytes (Balakrishnan et al., 2009). THAP1 plays a role in apoptosis by facilitating programmed cell death with the help of the transcription repressor protein Par-4 (Roussigne et al., 2003). LRRC49/THAP10 bidirectional gene pair is reported to have reduced expression in breast cancer (De Souza Santos et al., 2008). THAP11 is differentially expressed during human colon cancer progression and acts as a cell growth suppressor by negatively regulating the c-Myc pathway in gastric cancer (Zhang et al., 2020).

THAP9-AS1 (THAP9 antisense) is a newly annotated lncRNA gene encodes for long non-coding RNAs (Dreos et al., 2015; Howe et al., 2021). THAP9-AS1 lncRNA has been implicated in pancreatic cancer (N. Li et al., 2020), septic shock (Wu et al., 2020), and gastric cancer (Jia et al., 2019). It has also been reported recently that THAP9 and THAP9-AS1 exhibit different gene expression patterns under different types of stresses in the S-phase of the cell cycle. THAP9-AS1 is consistently upregulated under stress, whereas THAP9 exhibits downregulation and upregulation in other stress conditions. Both THAP9 and THAP9-AS1 exhibit a periodic gene expression throughout the S-phase, which is a characteristic of cell cycle-regulated genes (Sharma et al., 2021). Nevertheless, little is known about the biochemical and biological functions of the proteins encoded by THAP9 and THAP9-AS1 or their combined role in tumorigenesis.

In this study, we were interested in understanding the role of THAP9 and THAP9-AS1 in various cancers. We conducted a pan-cancer analysis of THAP9 and THAP9-AS1 expression, patient prognoses, and genetic mutations in TCGA (Tomczak et al., 2015) and GTEx datasets (GTEx Consortium, 2013) via TIMER2 (T. Li et al., 2020), GEPIA2 (Tang et al., 2019), EdgeR (Robinson et al., 2010) and cBio portal (Cerami et al., 2012). Moreover, we used Weighted gene co-expression network analysis (WGCNA) (Langfelder and Horvath, 2008) and Differential Gene correlation analysis (DGCA) (McKenzie et al., 2016) to explore the RNA-Seq datasets from TCGA for correlations between THAP9 and THAP9-AS1 genes and their correlation with other genes. Gene ontology (GO) and KEGG pathway enrichment analysis were performed to identify the primary biological functions linked to the genes that share the THAP9 and THA9-AS1 clusters in various tumor and normal samples. Our findings indicate statistical correlations between the expression of THAP9 and THAP9-AS1 with clinical prognosis, genetic mutations, and several cancer-related pathways, which suggests that they can serve as potential prognostic biomarkers for cancers.

## Results

### 1. Characterization of THAP9/THAP9-AS1 Promoter

THAP9 is a domesticated transposase (Majumdar et al., 2013, p. 9), and THAP9-AS1 (THAP9-anti sense1) is a lncRNA (Howe et al., 2021). Together they form a sense and antisense gene pair organized in a “head-to-head” orientation (**Figure 1a**). The sequence between an H2H gene pair, i.e., intra-H2H pair, can act as a bidirectional promoter (Li et al., 2006).

**Figure 1:**
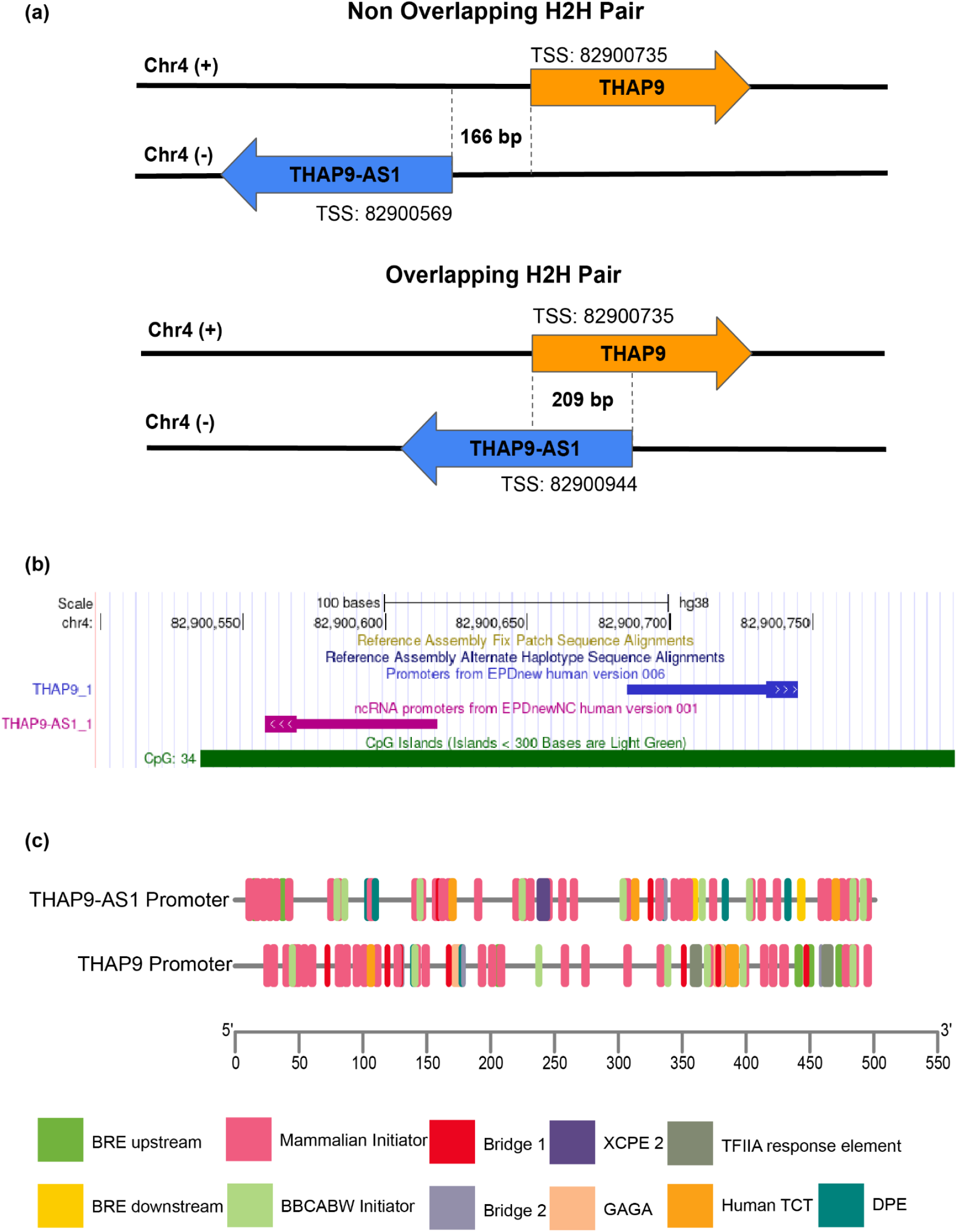
Identification and characterization of bidirectional THAP9/THAP9-AS1 promoter. **(a)** Schematic representation of the bidirectional genomic organization of THAP9 and THAP9-AS1 genes along with the TSS predicted by EPDnew. **(b)** UCSC genome browser showing THAP9 and THAP9-AS1 genes transcribed divergently based on the human GRCh38 assembly. CpG islands overlapping with the bidirectional promoter region are also indicated. **(c)** Schematic representation of the core promoter elements predicted by ElemeNT. The core promoter sequence used is −250 to +250 relative to the TSS of THAP9 and −400 to +100 for THAP9-AS1 (considering 82900569 as TSS) from EPDnew. The diagram is roughly to scale and constructed using TBtools.

Here we performed an *in silico* analysis of the putative THAP9/THAP9-AS1 bidirectional promoter region, that is the genomic region spanning the predicted THAP9/THAP9-AS1 transcriptional start sites (TSS). According to EPDnew (Dreos et al., 2015), the TSS of THAP9 is located at position 82900735 (**Supplementary Figure 1a**) on the sense strand of chromosome 4, while two TSSs were predicted for THAP9-AS1 at positions 82900569 and 82900944 (**Supplementary Figure 1b and c**), both located on the antisense strand of chromosome 4. Thus the predicted intergenic region between THAP9 and THAP9-AS1 is 166 bp (non-overlapping if THAP9-AS1 TSS is at position 82900569) or 209 bp (overlapping if THAP9-AS1 TSS is at position 82900944). In both cases, the THAP9-THAP9-AS1 sense antisense pair follows the head-to-head architecture (**Figure 1a**). For further characterization of the predicted bidirectional promoter region, we downloaded the sequence −250 to +250 relative to the predicted THAP9 TSS and −400 to +100 assuming position 82900569 as TSS (the selected sequence includes both TSSs for THAP9-AS1) from EPDnew.

The presence of CpG islands characterizes most bidirectional promoters. These regions are primarily devoid of DNA methylation and have a higher G + C content (Antequera, 2003). Thus, we examined the presence of CpG islands (CGI) in the THAP9/THAP9-AS1 bidirectional promoter region using the downloaded promoter sequences. Analysis and visualization of the selected promoter sequences for each gene, i.e., THAP9 and THAP9-AS1 using EMBOSS CPGplot (https://www.ebi.ac.uk/Tools/seqstats/emboss_cpgplot/), established that they individually fulfilled the criteria of CGI (Gardiner-Garden and Frommer, 1987) and showed GC content (Percent C + Percent G) > 50.00 and observed/expected CpG ratio > 0.60 (**Supplementary Figure 1c and d**). We then looked for already annotated CGIs around the THAP9/THAP-AS1 bidirectional promoter region using the UCSC Genome Browser (http://genome.ucsc.edu/). We observed a CGI located between positions 82900535 and 82900912 on chromosome 4 overlapping with the THAP9/THAP-AS1 promoter (**Figure 1b**). It is tempting to speculate that differential DNA methylation of this CGI may influence bidirectional gene expression of THAP9/THAP-AS1.

In addition to CGI, bidirectional promoters are often enriched with specific histone marks. ENCODE (Encyclopedia of DNA Elements) (Davis et al., 2018) (EH38E3592191, EH38E3592192) data revealed the presence of bimodal peaks of transcriptionally active histone modifications at the THAP9/THAP9-AS1 promoter region; this strongly suggests bidirectional transcriptional activity (**Figure 2**) (Bornelöv et al., 2015). Moreover, the THAP9/THAP9-AS1 bidirectional promoter was also characterized by DNase I hypersensitivity.

**Figure 2:**
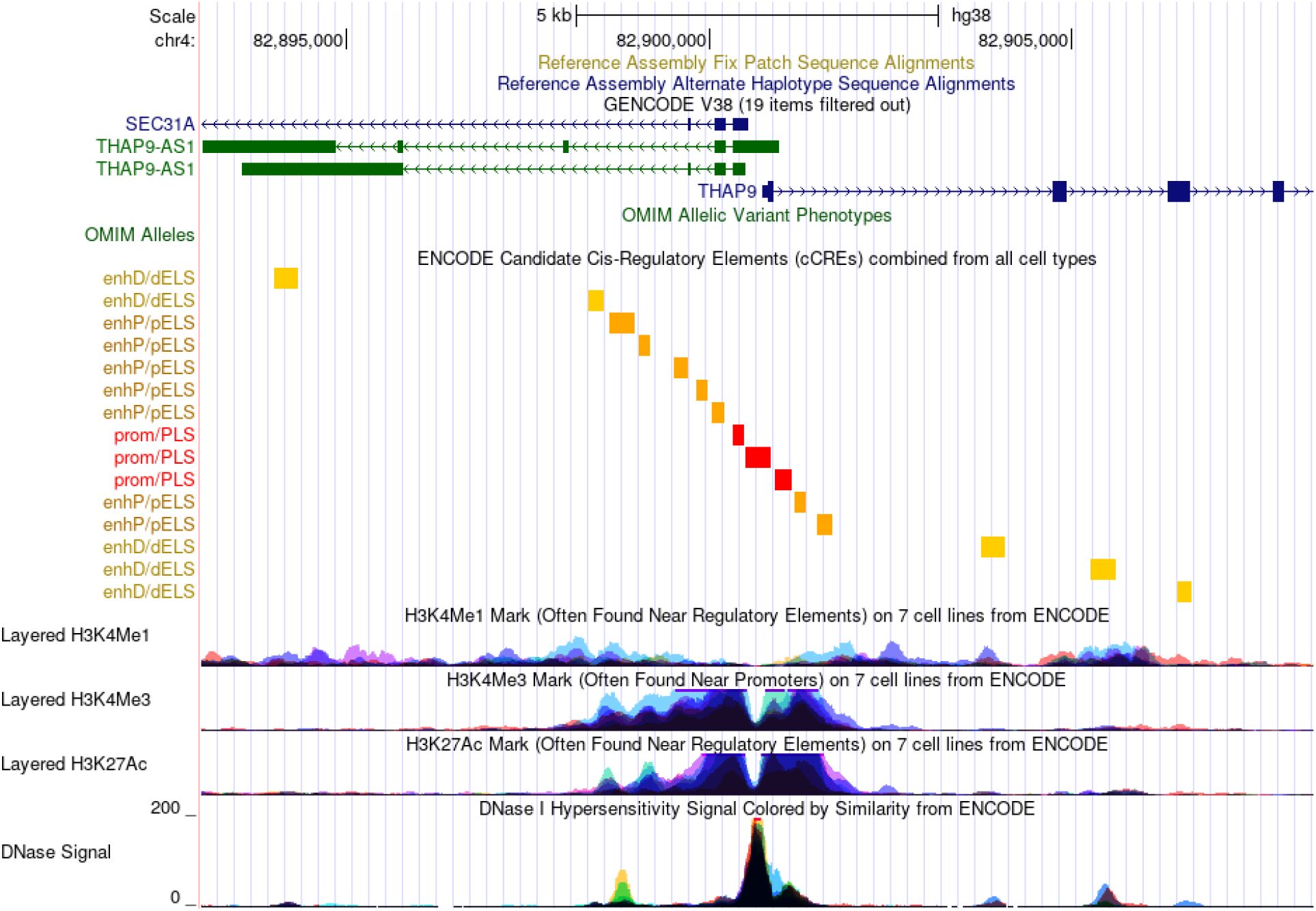
UCSC Genome Browser representing ENCODE data for THAP9/THAP9-AS1 Bidirectional Promoter Region. The genomic region contains the putative bidirectional promoter region of the THAP9/THAP9-AS1 gene pair. The GENCODE Genes track shows transcript variants for both genes. Below that is the ENCODE Candidate cis-Regulatory Elements (cCREs) track, which shows the presence of several regulatory elements in the promoter region. The next three tracks are from the ENCODE project showing the H3K4Me1 mark, H3K4me3 mark, and the H3K27Ac mark associated with active regulatory regions in 7 cell lines. Moreover, the peak in the DNAse I Hypersensitivity Signal showed in the last track also indicates a large number of regulatory regions.

While there is no consensus computational method to determine whether a promoter is bidirectional or not, certain core promoter elements are known to be essential structural features of bidirectional promoters. These elements include the TATA box, CCAAT box, B recognition element (BRE), initiator element (INR), and downstream promoter element (DPE) (Yang and Elnitski, 2008). The TATA box exists in both unidirectional and bidirectional promoters; however, they are less frequent amongst bidirectional promoters. Bidirectional promoters generally have a higher enrichment of CCAAT boxes and the BRE element than unidirectional promoters, while the ratio of DPE and INR remains unchanged mainly (Yang and Elnitski, 2008).

Therefore we submitted the THAP9 and THAP9-AS1 promoter sequences to ElemeNT (https://www.juven-gershonlab.org/resources/element/run/) (Sloutskin et al., 2015) to predict putative core promoter elements involved in the regulation of THAP9/THAP9-AS1 gene pair (**Figure 1c**). Potential regulatory elements identified within the THAP9 and THAP9-AS1 promoters include BRE, GAGA, Mammalian Initiator, Bridge 1, DPE, BBCABW Initiator, Human TCT, TFIIA response element, but no TATA-Box (**Figure 1c, Supplementary Table 1, 2**). In the absence of a TATA box, the DPE is required for binding the transcription factor TFIID, and DPE acts together with INR (Smale and Kadonaga, 2003). DPE is primarily recognized by TBP associated factors (TAFs), particularly TAF6 and TAF9 (Kutach and Kadonaga, 2000). The INR element signifies the point of transcription initiation in the TATA-less promoter. INR is functionally similar to the TATA box, but it can function independently (Yang et al., 2007). Moreover, in the TATA-less promoters, TFIIB–BRE interaction plays a vital role in assembling the preinitiation complex and transcription initiation (Lagrange et al., 1998). Thus, we can say that the selected THAP9/THAP9-AS1 bidirectional promoter displays several characteristics of bidirectional promoters.

### 2. Difference Between THAP9/THAP9-AS1 Expression in several cancers

To compare the expression levels of THAP9 and THAP9-AS1 genes between tumor and normal samples, we analyzed their expression across various cancer types using the TCGA (Tomczak et al., 2015) and GTEx (GTEx Consortium, 2013) datasets via EdgeR (Robinson et al., 2010), TIMER2.0 (T. Li et al., 2020, p. 2) and GEPIA2 (Tang et al., 2019, p. 2).

#### TIMER2

We first used TIMER2.0 to evaluate the expression of THAP9 between primary tumor and normal samples using the TCGA database. We found that THAP9 expression levels in tumor tissue of CHOL (P < 0.001), COAD (P < 0.001), ESCA (P < 0.01), LIHC (P < 0.001), LUSC (P < 0.001), LUAD (P < 0.05), and STAD (P < 0.001) are higher than the corresponding normal tissue based on the TCGA dataset (**Figure 3a**). On the contrary, THAP9 expression levels in tumor tissue KIRC (P < 0.001), KIRP (P < 0.001), PRAD (P < 0.05), THCA (P < 0.001), and UCEC (P < 0.001) are lower than the corresponding normal tissue (**Figure 3a**). We could not perform a similar comparison in ACC, DLBC, LAML, LGG, MESO, OV, SARC, SKCM, TGCT, UCS, and UVM because they lack normal samples in the TCGA database. A similar analysis could not be conducted for THAP9-AS1 since TIMER2.0 does not give any gene expression profile for THAP9-AS1.

**Figure 3:**
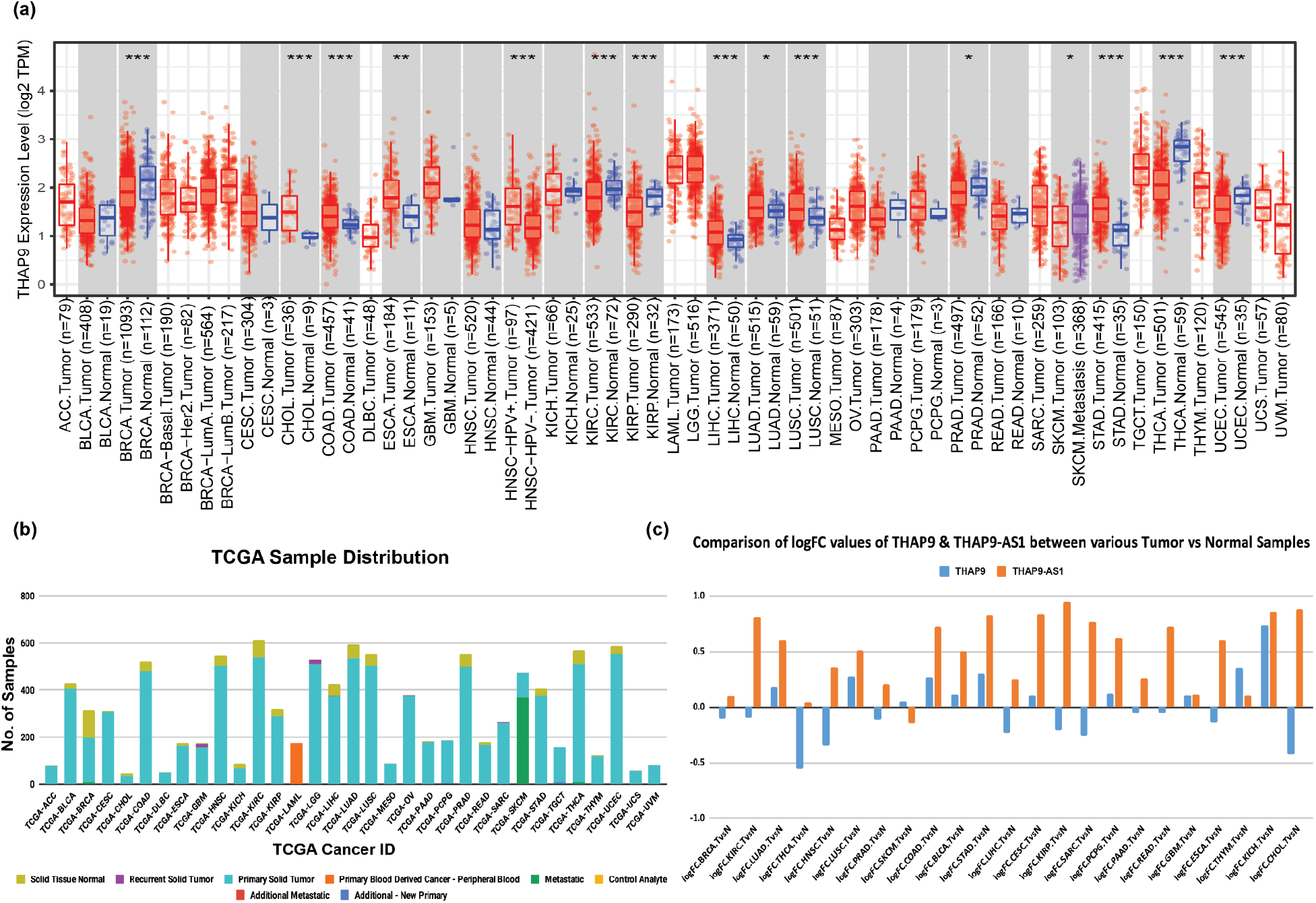
THAP9 and THAP9-AS1 gene expression levels in different tumors. **(a)** THAP9 expression levels in human tumors (red) and corresponding normal tissues (blue) were obtained through TIMER2. **(b)** Distribution of RNA-Seq HT-Seq counts datasets downloaded from TCGA (**Supplementary Table 3**) **(c)** The logFC values of THAP9 (blue) and THAP9-AS1 (orange) after differential gene expression analysis using EdgeR (**Supplementary Table 4**).

#### EdgeR

We then compared the gene expression profiles of 10,490 samples from 23 TCGA (**Supplementary Table 3**) cancer types between tumor (primary solid tumor) and normal (solid tissue normal) samples. By comparing the TCGA HTSeq read counts (**Figure 3b**) of 34125 genes using EdgeR, it appeared that THAP9 and THAP9-AS1 genes were not differentially expressed in any of the cancers, with their log2FC not above 1 in any of the normal vs. tumor pairs (**Figure 3c, Supplementary Table 4**).

#### GEPIA2

The TIMER2.0 database did not have THAP9-AS1 data and TCGA did not have normal samples for ACC, DLBC, LAML, LGG, MESO, OV, SARC, SKCM, TGCT, THYM, UCS, and UVM. Thus, to get a more broad understanding of the gene expression profiles of the two genes we used GEPIA2, which analyzes genes using both the TCGA and GTEx databases. It helped us obtain the differential gene expression profiles of THAP9 and THAP9-AS1 across all 33 cancers. We observed that THAP9 expression was downregulated in TGCT (P <0.01) and was upregulated in THYM (P < 0.01), (**Figure 4a, b**). However, THAP9-AS1 showed downregulation in OV (P < 0.01), SKCM (P < 0.01), and THCA (P < 0.01) (**Figure 4c,d, e**) and was upregulated in THYM (P < 0.01), PAAD (P < 0.01), DLBC (P < 0.01), and CHOL (P < 0.01) (**Figure 4f,g,h,i**). Moreover, from the above results, we observed that both the genes are simultaneously downregulated in THYM. Thus, the differential expression of THAP9 and THAP9-AS1 in different tumor types suggests that the two genes have diverse regulatory mechanisms in different tumor types.

**Figure 4:**
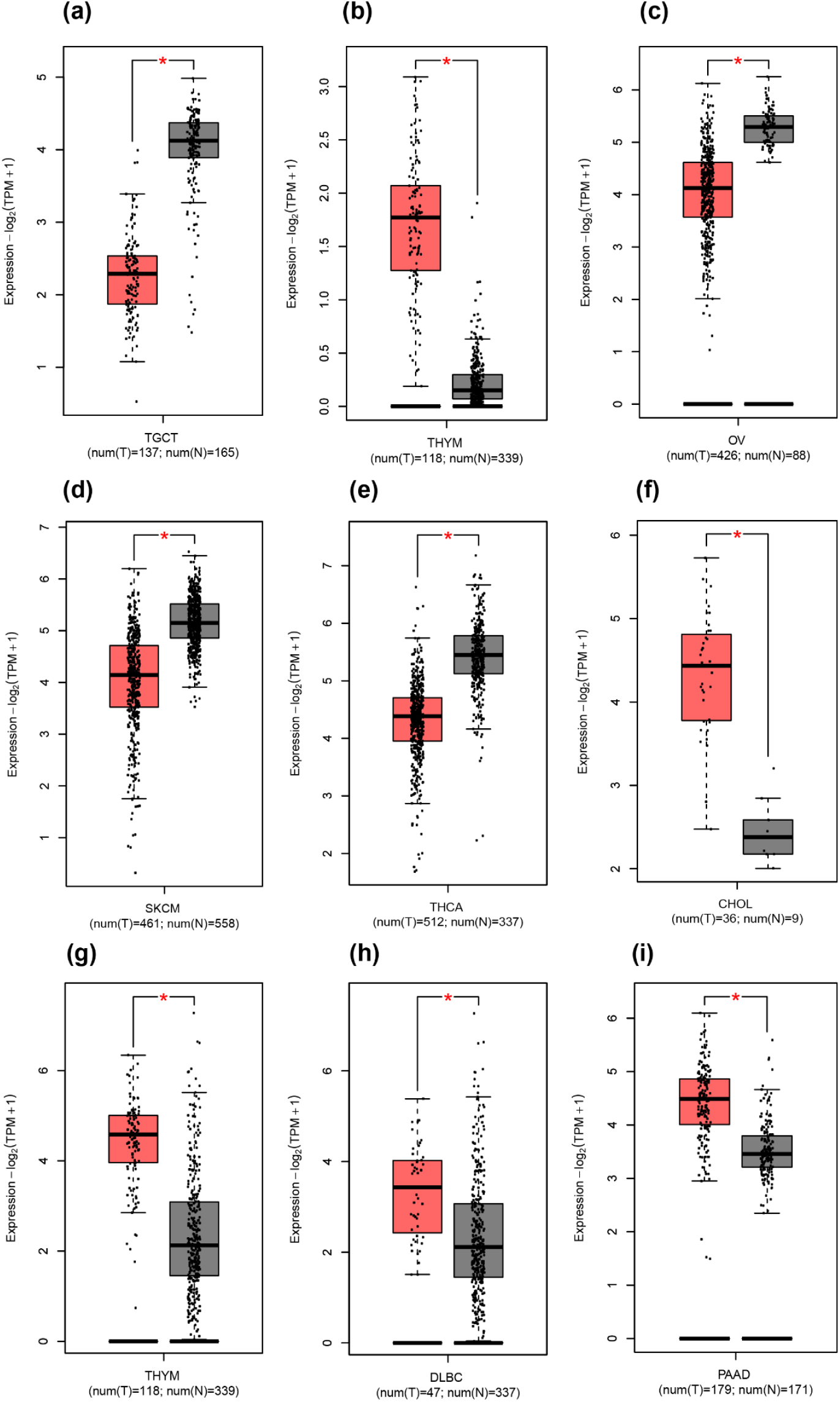
Box plot representation of the comparative expression of THAP9 (a) and (b) and THAP9-AS1 (c)-(i) in different tumor samples (red) vs. normal tissue samples (grey) from TCGA and GTEx generated using GEPIA2. * P<0.01. GEPIA2 uses one-way ANOVA, using the pathological stage (X-axis) as the variable for performing differential expression of the input gene. The expression data used for the analysis is log2(TPM+1) (y-axis) transformed. **(a)** THAP9 is downregulated in TGCT and **(b)** upregulated THYM. **(c) - (e)** THAP9-AS1 is downregulated in OV, SKCM, and THCA, and **(f) - (i)** it is upregulated in CHOL, THYM, DLBC, and PAAD. (*P < 0.05).

### 3. The prognostic analysis of THAP9 and THAP9-AS1

We used the dataset from TCGA and GTEx via GEPIA2 to investigate the correlation of THAP9 and THAP9-AS1 expression with patients’ prognoses across different tumor types. The Survival heat map of hazard ratio (HR) for overall and disease-free survival (**Figure 5a**, **6a**) shows the prognostic impacts of THAP9 and THAP9-AS1 in multiple cancer types. Poor prognosis and overall survival were linked to upregulation of THAP9 expression in LGG and STAD and its downregulation in HNSC and KIRC (**Figure 5b**). Moreover, poor DFS (Disease-Free Survival) was linked with upregulated THAP9 expression in BLCA and CESC and its downregulated expression in KIRC and THYM (**Figure 6b**). Similarly, in the case of THAP9-AS1, its upregulation was linked to poor prognosis and overall survival in ACC, LGG, PRAD, SARC, and THCA (**Figure 5c**) and poor DFS in ACC, KICH, and MESO, while its down-regulated expression was linked to poor DFS in KIRC (**Figure 6c**).

**Figure 5:**
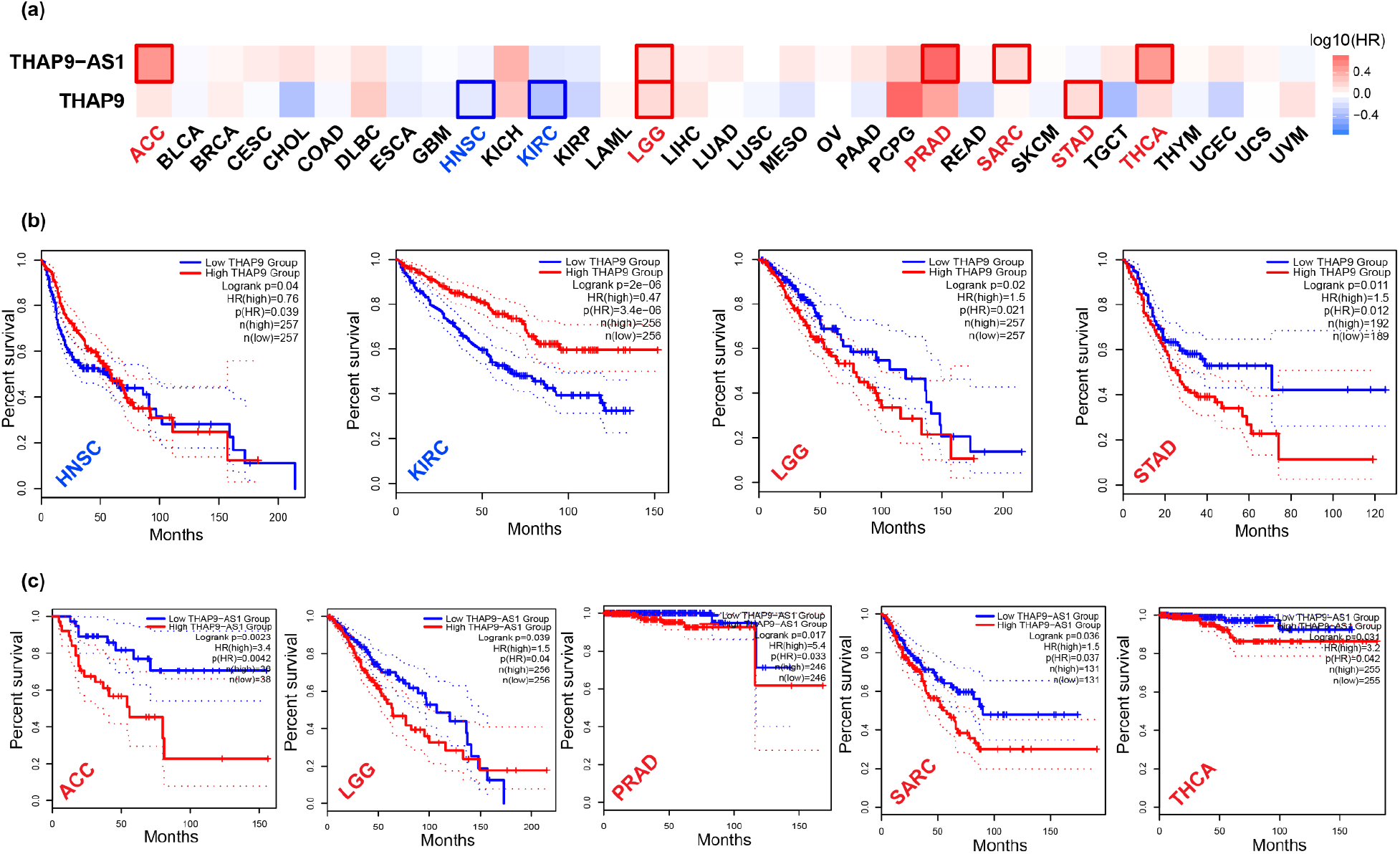
Overall patient survival analysis using GEPIA2. **(a)** The relationship between THAP9 and THAP9-AS1 gene expression and the overall survival prognosis of cancers in TCGA. Median is selected as a threshold for separating high-expression and low expression cohorts. The red and blue blocks represent higher and lower risks, respectively. The bounding boxes depicted the significant (p < 0.05) unfavorable and favorable results, respectively. The overall survival rate and gene expression (from TCGA) of **(b)** THAP9 in HNSC, KIRC, LGG, and STAD **(c)** THAP9-AS1 in ACC, LGG, PRAD, SARC, and THCA.

**Figure 6:**
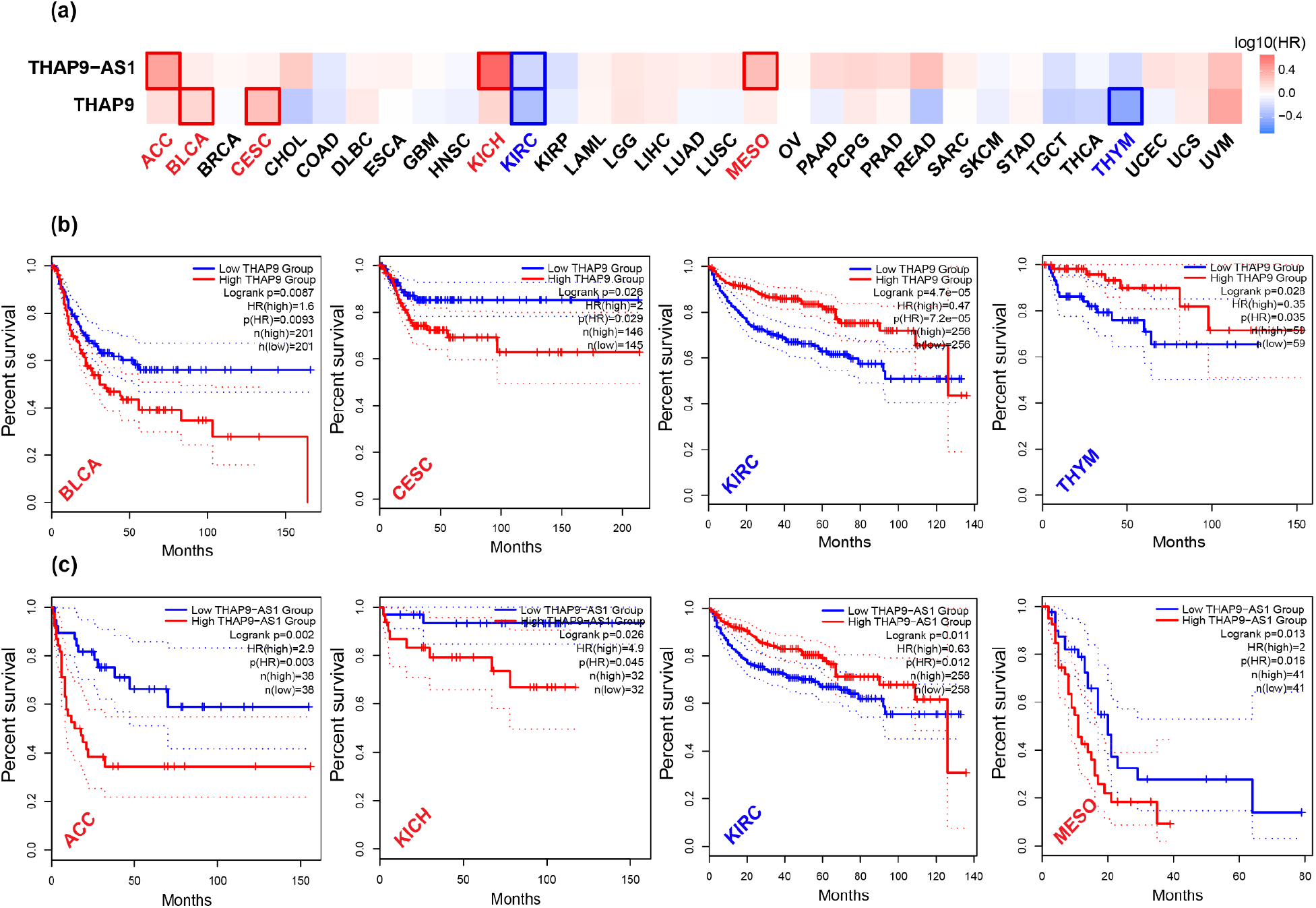
Disease Free survival analysis using GEPIA2. **(a)** The relationship between THAP9 and THAP9-AS1 gene expression and the Disease-Free Survival prognosis of cancers in TCGA. Median is selected as a threshold for separating high-expression and low expression cohorts. The red and blue blocks represent higher and lower risks, respectively. The bounding boxes depicted the significant (p < 0.05) unfavorable and favorable results, respectively. The Disease-Free survival rate and gene expression of **(b)** THAP9 in BLCA, CESC, KIRC, and THYM **(c)** THAP9-AS1 in ACC, KICH, KIRC, and MESO.

### 4. Analysis of THAP9/THAP9-AS1 mutations in various tumors

To study the prevalence of mutations in the THAP9 and THAP9-AS1 genes across various human cancers, we used the cBioPortal tool (Cerami et al., 2012) in the “Pan-cancer analysis of whole genomes (ICGC/TCGA, Nature 2020)” dataset available at http://www.cbioportal.org. As shown in **Figures 7(a) and (d)**, we found that pancreatic cancer patients had the highest alteration frequency in both THAP9 and THAP9-AS1 (> 6%) with “amplification” (i.e., more copies, often focal) as the primary alteration type. Notably, all patients with breast cancer, Non-Small Cell Lung Cancer, melanoma, embryonal tumor, and bone cancer also had amplification of both THAP9 and THAP9-AS1 genes, and all the patients of Uterine Endometrioid Carcinoma showed “deep deletion” (indicates a deep loss, possibly a homozygous deletion) of THAP9 and THAP9-AS1 genes. Moreover, for the THAP9 gene, “mutation” appeared as the only form of genetic alteration in all patients with colorectal cancer, mature B-cell Lymphoma, and Head and Neck Cancer. The sites and types of THAP9 genetic alteration are presented in **Figure 7c (Supplementary Table 5)**. The median months’ survivals were 37.73 and 49.61 for the altered THAP9 group (**Figure 7b**) and the reference group respectively and *NA* and 49.61, respectively for the altered THAP9-AS1 group (**Figure 7e**) and the reference group. Thus, the clinical survival prognosis value of THAP9 and THAP9-AS1 alterations reflected poor overall survival in the altered group.

**Figure 7:**
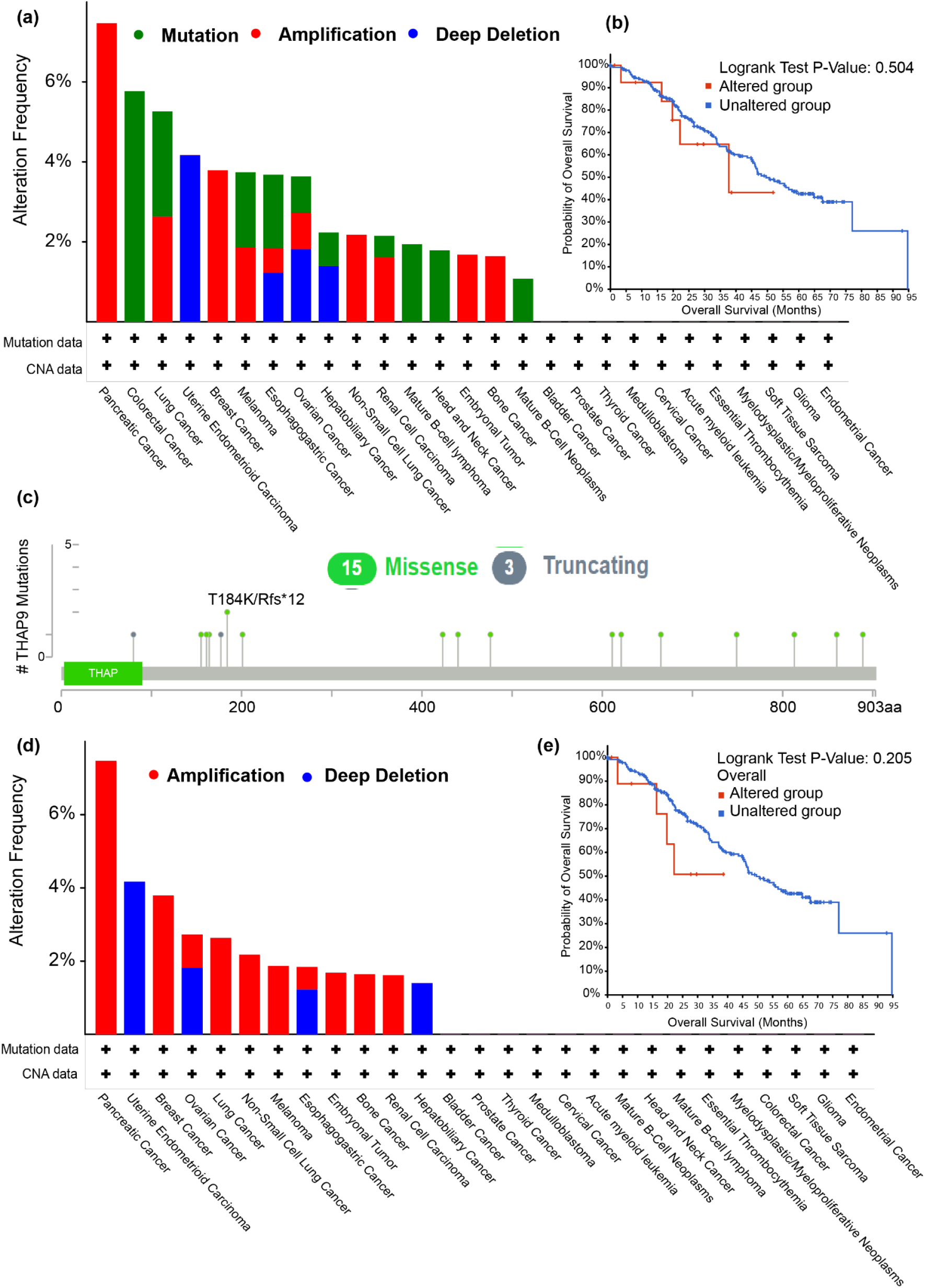
Mutation of THAP9 and THAP9-AS1 in different cancers in TCGA. The alteration frequencies with mutation type for **(a)** THAP9 **(d)** THAP9-AS1, where X-axis represents the type of alteration (red-amplification, blue - deep deletion, and green - mutation) and Y-axis represents the frequency of alteration in different cancers. Correlation between mutation status and overall survival of cancer patients in **(b)** THAP9 **(e)** THAP9-AS1. The red line shows the overall survival estimate for patients with alteration in the gene as compared to patients with no alteration (blue line). Survival analysis significance was based on the Log-rank Test. P < 0.05 was considered significant. **(c)** Mutation sites in THAP9 (Refer to Supplementary Table for Details).

### 5. Understanding the role of THAP9 and THAP9-AS1 using guilt-by-association analysis

GBA (Guilt By Association) (Oliver, 2000) analysis is often used to predict an unknown gene’s function by grouping it with known genes that share its transcriptional behavior. Genes turned on or turned off together under various conditions may be part of the same cellular processes (van Dam et al., 2017). The exact cellular functions of THAP9 and THAP9-AS1 are unknown. Thus, we decided to compare the expression of THAP9 with THAP9-AS1 and 34125 other genes from over 9571 patients across 22 human cancers (and associated normal samples) fetched from TCGA. The gene co-expression network for THAP9 and THAP9-AS1 was constructed using WGCNA followed by a differential gene correlation analysis using DGCA. We also investigated Gene Ontology and KEGG pathways for the genes coexpressed with the two genes and the genes differentially correlated across normal vs. tumor samples. The functions of the coexpressed genes should provide more insights into the physiological functions as well as the possible role of THAP9 and THAP9-AS1 in cancers.

#### 5.1 Gene coexpression analysis

The first step of GBA is to identify the genes that are co-expressed with THAP9 and THAP9-AS1 in normal vs. tumor samples in each cancer type. We utilized the WGCNA R package (Langfelder and Horvath, 2008) to build a weighted co-expression network for THAP9 and THAP9-AS1. The samples of each tumor and normal pair (considered cancers with more than 3 paired normal samples) were clustered to identify the gene modules representing genes co-expressed with THAP9 and THAP9-AS1. We selected the top 20 genes coexpressed with THAP9 and THAP9-AS1 in each condition to identify the frequently co-expressed genes (**Figure 8, Supplementary Data available on Github**).

**Figure 8:**
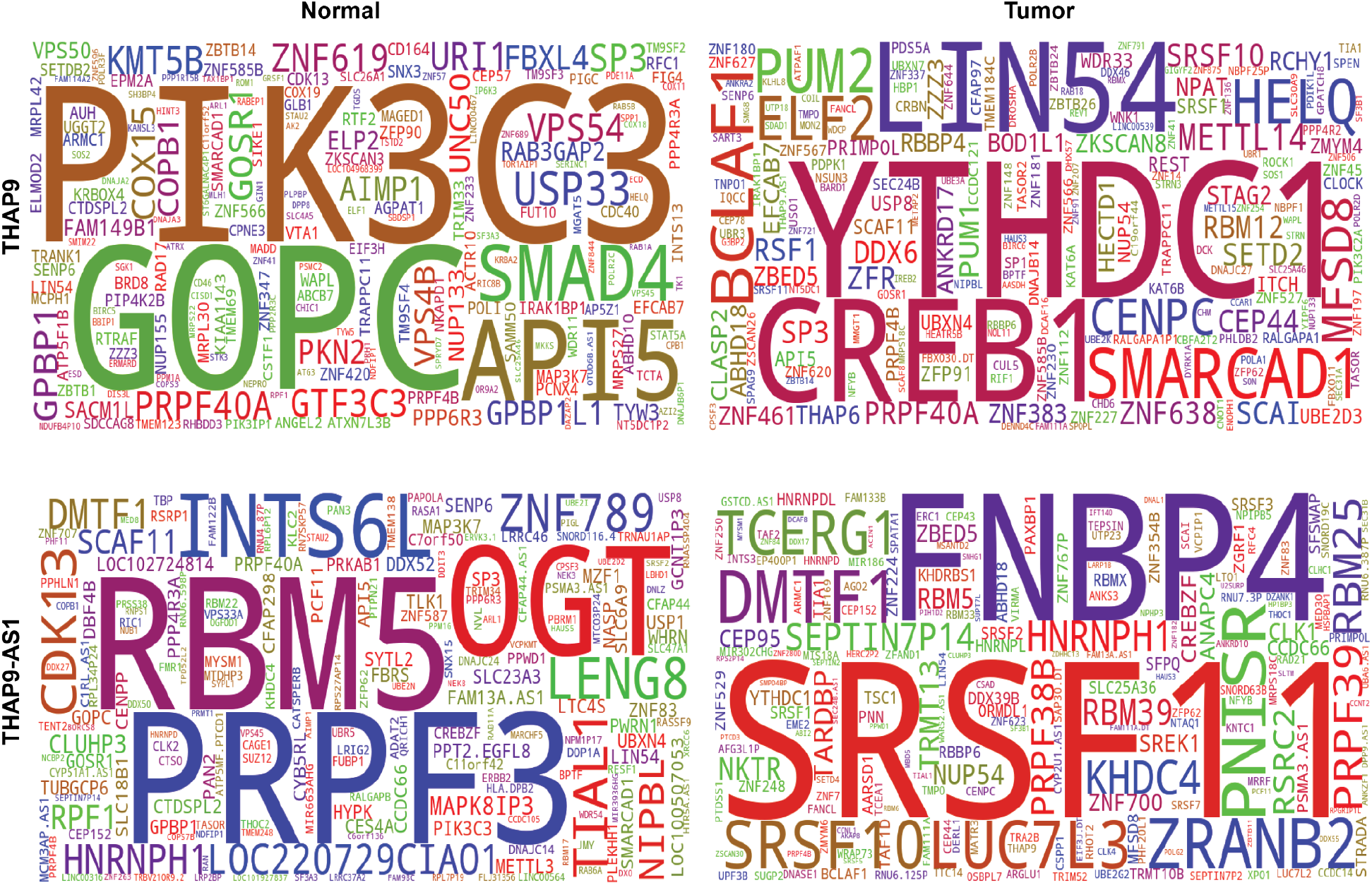
Consensus of genes coexpressed with THAP9 and THAP9-AS1. Word Cloud of top 20 genes frequently co-expressed with: (1st row) THAP9 in different normal (top-left) vs. tumor (top-right) samples. (2nd row) THAP9-AS1 in different normal (bottom-left) vs. tumor (bottom-right) samples.The coexpressing genes are identified using the WGCNA Bioconductor package and plotted using the Wordcloud python package. The height of the word is directly proportional to the frequency of coexpression with THAP9.

The second step is to identify the functional association of the THAP9 and THAP9-AS1 coexpression modules. It will help us get a deeper insight into the physiological role of THAP9 and THAP9-AS1 in each condition. We used the ‘ShinyGO,’ a Shiny application developed based on several R/Bioconductor packages, which performs in-depth analysis of gene lists, which includes graphical visualization of enrichment, pathway, gene characteristics, and protein interactions (Ge et al., 2020). The noteworthy pathways from GO analysis for THAP9 and THAP9-AS1 were visualized in **Figure 9** and **Figure 10**, respectively (**Supplementary Table 6 and 7**).

**Figure 9:**
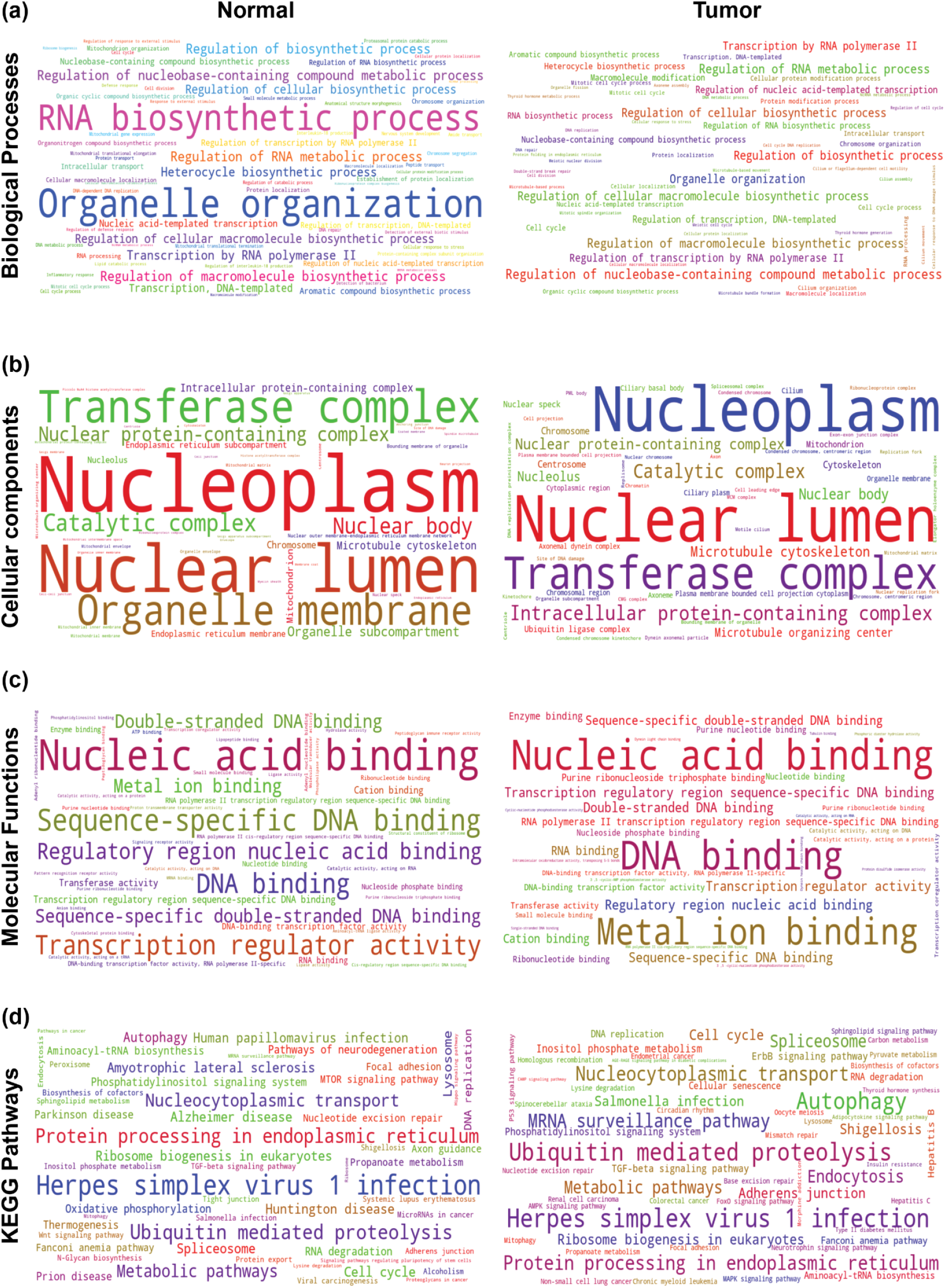
Gene ontology (GO) and KEGG pathway analysis of genes coexpressed with THAP9 in normal vs. tumor samples. The enrichment test was performed for normal vs. tumor samples (for each cancer) using the ShinyGO for the top 10 enriched terms, with the significance cutoff for adjusted p-value was set at 0.05. The font sizes in the word cloud are proportional to their frequency after the enrichment is merged for all the cancers (left-side for normal samples and right-side for tumor samples). **(a)** Word cloud of enriched GO terms in the biological process category. **(b)** Word cloud of enriched GO terms in the cellular component category. **(c)** Word cloud of enriched GO terms in the molecular functions category. **(d)** Word cloud of enriched KEGG Pathways.

**Figure 10:**
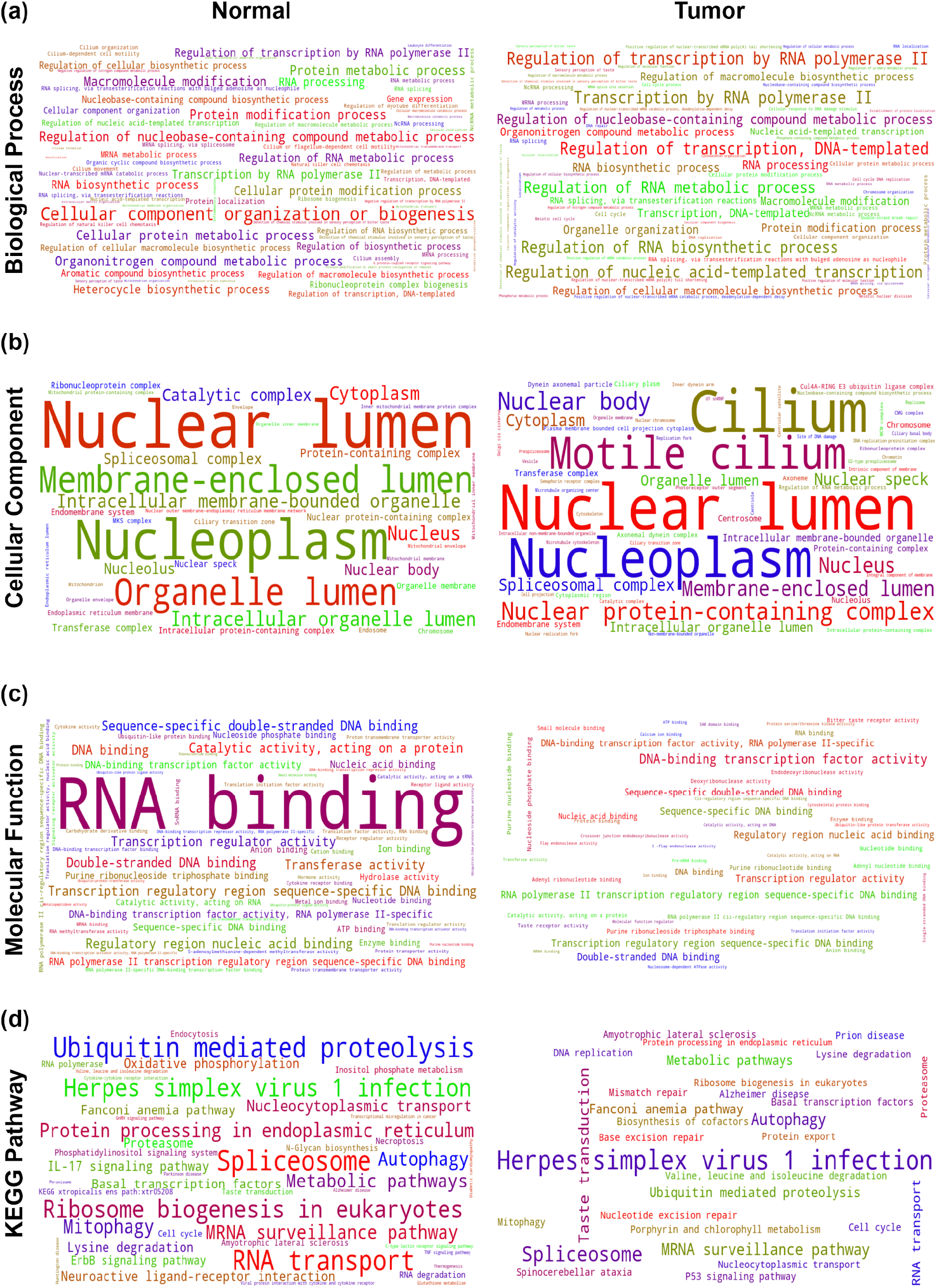
Gene ontology (GO) and KEGG pathway analysis of genes coexpressed with THAP9-AS1 in normal vs. tumor samples. The enrichment test was performed for normal vs. tumor samples (for each cancer) using the ShinyGO for the top 10 enriched terms, with the significance cutoff for adjusted p-value was set at 0.05. The font sizes in the word cloud are proportional to their frequency after the enrichment is merged for all the cancers (left-side for normal samples and right-side for tumor samples). **(a)** Word cloud of enriched GO terms in the biological process category. **(b)** Word cloud of enriched GO terms in the cellular component category. **(c)** Word cloud of enriched GO terms in the molecular functions category. **(d)** Word cloud of enriched KEGG Pathways.

##### THAP9

As presented in **Figure 9 (Supplementary Table 6)**, in the GO-BP (Biological Process) group, genes coexpressed with THAP9 in normal condition were primarily involved in RNA Biosynthetic process, Organelle organization, and more, while in tumor condition we observed some regulatory processes like Regulation of nucleobase-containing compound metabolic process, regulation of cellular biosynthetic process, and more. In the GO-CC (Cellular Component) group, the co-expressed genes in the normal and tumor samples were markedly enriched in Transferase complex, Nucleoplasm, Nuclear Lumen, and catalytic complex. In comparison, the enrichment of the Organelle membrane reduced between normal vs. tumor samples. In the GO-MF (Molecular Function) group, the significantly enriched pathways in normal samples were Nucleic acid-binding, Sequence-specific DNA binding, Regulatory region nucleic acid binding, DNA binding, and transcription regulator activity. In contrast, in tumor samples, the enriched MF pathways are Nucleic acid-binding, DNA binding, and Metal ion binding. According to KEGG pathway analysis, the most markedly enriched pathways in normal and tumor samples included Herpes simplex virus 1 infection, Protein processing endoplasmic reticulum, ubiquitin-mediated. Proteolysis, Nucleocytoplasmic transport, and Autophagy. Apart from that, several KEGG pathways associated with neurodegenerative disorders like Alzheimer’s disease, Parkinson’s disease, Huntington disease, and pathways for neurodegeneration were also enriched in the normal samples (**Figure 9**).

##### THAP9-AS1

Similarly, **Figure 10 (Supplementary Table 7)** shows that, in the GO-BP group, genes coexpressed with THAP9-AS1 in normal condition were primarily involved in Cellular component organization or biogenesis, while in tumor condition we observed some regulatory processes like Regulation of transcription by RNA polymerase II, Transcription by RNA polymerase II, Regulation of transcription, DNA templated, and more. Nevertheless, none of the pathways appeared to be significantly enriched in both cases. The co-expressed genes in the normal and tumor samples were markedly enriched in Nucleoplasm, nuclear lumen, and membrane-enclosed lumen in the GO-CC group. In contrast, the significant enrichment in Cilium and Motile Cilium appeared only in the tumor samples. The significantly enriched pathways in normal samples in the GO-MF group were RNA binding. Moreover, Molecular functions like Sequence-specific double-stranded DNA binding, Regulatory region nucleic acid binding, transcription regulator activity, RNA polymerase II transcription regulatory region sequence-specific DNA binding, etc appear enriched in normal and tumor samples. According to KEGG pathway analysis, the most notably enriched pathways in normal and tumor samples included Herpes simplex virus 1 infection, Spliceosome, and Ubiquitin mediated proteolysis, Nucleocytoplasmic transport, and Autophagy (**Figure 10**).

#### 5.2 Differential Gene Correlation analysis

Differential gene correlation/co-expression analysis can identify biologically important differentially correlated genes that can not be detected using regular Gene co-expression analysis or differential gene expression analyses. Literature suggests that differentially correlated genes between different sample groups are more likely to be the regulators and explain differences between phenotypes (Amar et al., 2013; Hudson et al., 2009; Kostka and Spang, 2004). Multiple studies have used differential correlation analyses to identify genes underlying differences between healthy and disease samples or between different tissues cell types or species (Gao et al., 2012; Monaco et al., 2015; Pierson et al., 2015; Zeisel et al., 2015). Considering that the genes that are functionally related tend to have similar expression profiles; therefore, differential gene correlation analysis that can compare the expression correlation of THAP9 and THAP9-AS1 with other genes in normal vs. tumor samples can give us insight into biological processes and molecular pathways that distinctly involves the two genes between the two conditions.

DGCA is an R package (McKenzie et al., 2016) designed to detect differences in the correlations of gene pairs between distinct biological conditions. DGCA uses correlation coefficients transformed into normalized Z-scores to identify differentially correlated genes and modules while performing downstream analysis, including data visualization, GO enrichment, and network construction tools. In this study, we used DGCA to identify the genes differentially correlated with THAP9 and THAP9-AS1 under various tumor vs. paired normal conditions.

We used the RNA-seq HT-Seq count dataset of 21 cancer and paired normal samples from TCGA. We restricted our analysis to identify the genes differentially correlated with THAP9 and THAP9-AS1 in the tumor vs. normal sample group. First we calculated the correlation between THAP9 and THAP9-AS1 expression between normal and tumor samples in each cancer type and they appeared positively correlated in each condition suggesting their coordinated regulation (**Supplementary Figure 4**). We calculated the difference in Spearman correlations for all the genes with THAP9 and THAP9-AS1 to identify the genes differentially correlated with THAP9 and THAP9-AS1 between the tumor and the paired normal sample in each cancer type. Then we measured the gene ontology (GO) enrichment of the genes differentially correlated with THAP9 and THAP9-AS1 with a gain and loss of correlation in tumor vs. normal samples.

##### THAP9

When we took consensus of Gene Ontology terms associated within the genes that gained or lost correlation with THAP9 in all the cancer samples (**Figure 11**, Result for each cancer separately in Supplementary Figure 2), we observed that pattern specification process, anatomical structure morphogenesis, nervous system development, etc. were some of the biological processes enriched in the genes that gained correlation with THAP9 while transposition, DNA mediated, biomineral tissue development, negative regulation of gene expression, etc. were the genes that lost correlation with THAP9. In the cellular component group, Golgi apparatus, lytic vacuole, extracellular matrix, etc., were the terms enriched among the genes that gained correlation with THAP9. In contrast, nuclear chromatin, extracellular space, supramolecular fiber, etc. are the terms enriched among the genes that lost correlation with THAP9. For molecular function group genes that gain correlation with THAP9 were enriched in catalytic activity, glycosaminoglycan binding, anion binding, etc., while the genes that lost correlation with THAP9 were enriched in cation channel activity, transposase activity, voltage-gated ion channel activity, etc.

**Figure 11:**
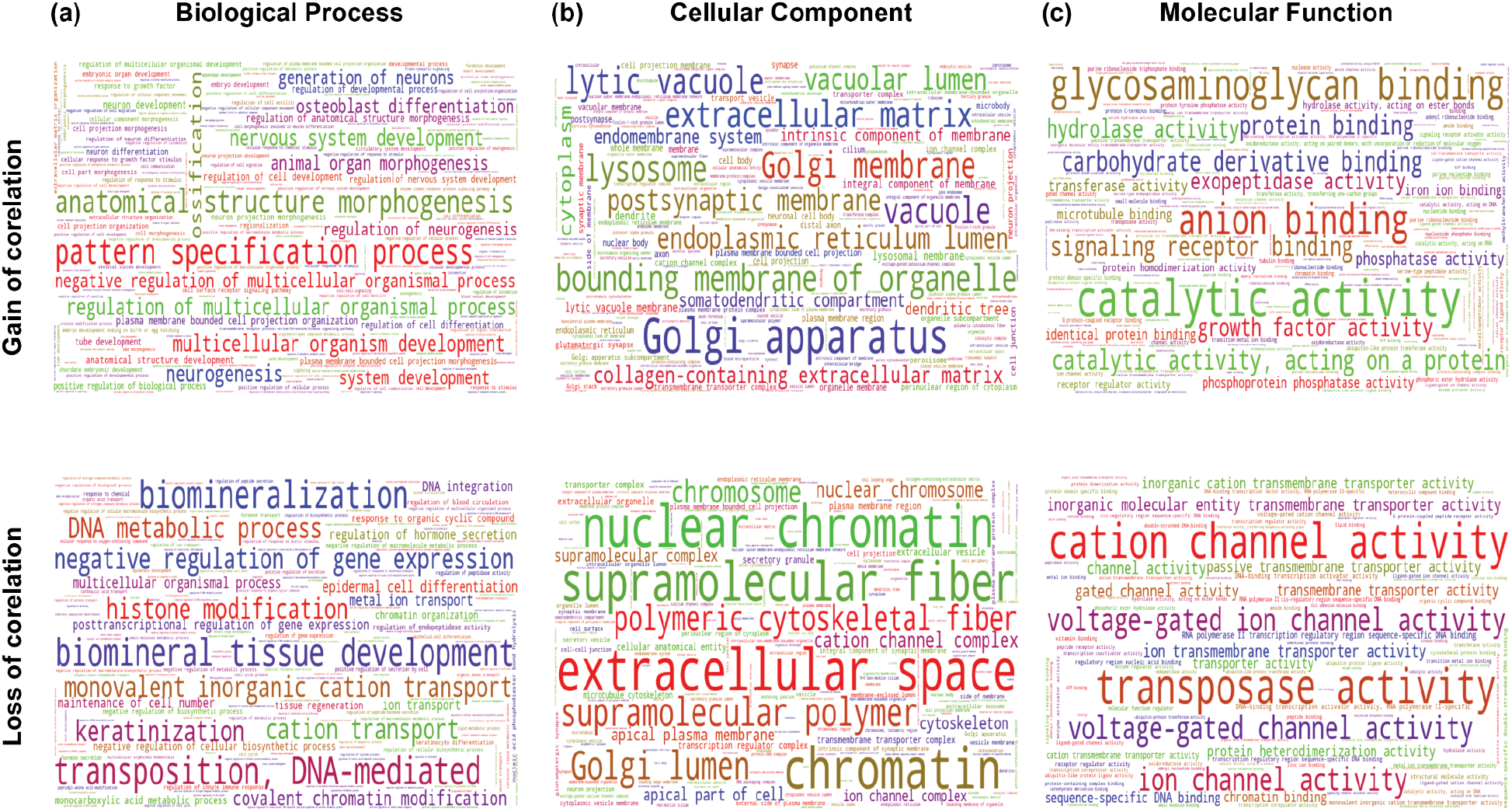
Gene Ontology Analysis of genes differentially correlated with THAP9 in Normal vs. Tumor samples. (Top - Genes that gained correlation with THAP9, Bottom - Genes that lost correlation with THAP9). Word cloud of Enriched **(a)** GO biological process **(b)** GO cellular component **(c)** GO molecular functions. The differential gene correlation analysis, followed by the gene ontology analysis for the differentially correlated genes, is performed using the DGCA R.

##### THAP9-AS1

*T*he analysis for THAP9-AS1 (**Figure 12**, Result for each cancer separately in Supplementary Figure 3) showed that the genes that gained correlation with THAP-AS1 were enriched in the immune system process, immune effector process, response to stimulus, leukocyte activation, cell activation, etc. were enriched BP group; vesicle, membrane, cytoplasm, endomembrane, etc. were enriched in CC group and catalytic activity, immune receptor activity, hydrolase activity, etc. were the terms enriched in MF group. On the other hand, genes that lost correlation with THAP9-AS1 were enriched in the developmental process, anatomical structure development, multicellular organismal process, multicellular organism development, etc. were processes enriched in the BP group: plasma membrane, cytoplasm, cell periphery, etc. were the terms enriched in the CC group and molecular function regulator, catalytic activity. Binding, transferase activity, protein binding, etc., were the terms enriched in the MF group.

**Figure 12:**
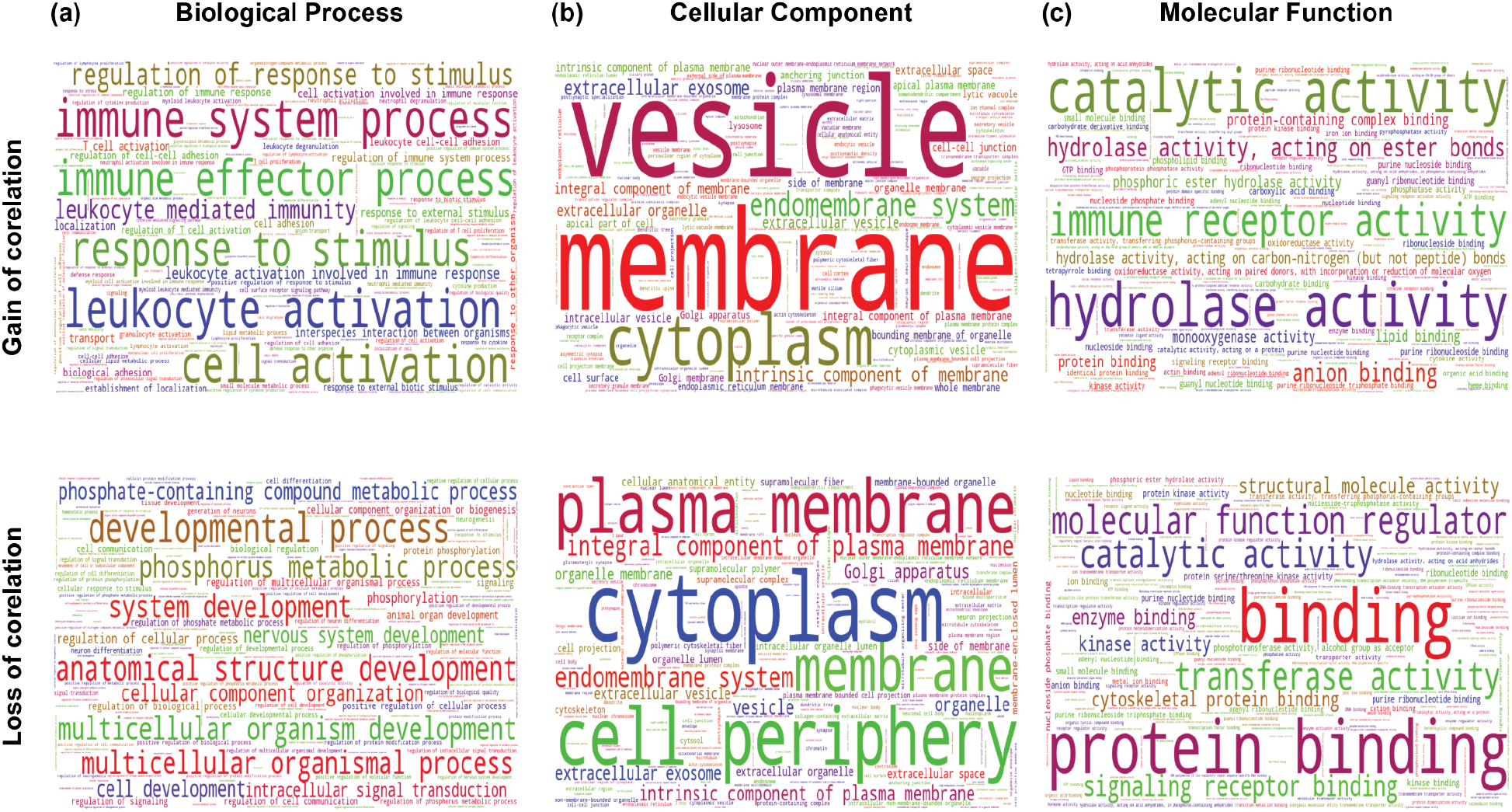
Gene Ontology Analysis of genes differentially correlated with THAP9 in Normal vs. Tumor samples. (Top - Genes that gained correlation with THAP9, Bottom - Genes that lost correlation with THAP9). Word cloud of Enriched **(a)** GO biological process **(b)** GO cellular component **(C)** GO molecular functions. The differential gene correlation analysis, followed by the gene ontology analysis for the differentially correlated genes, is performed using the DGCA R.

## Discussion

This study shows that THAP9 and THAP9-AS1 genes are located in a “head-to-head” orientation on chromosome 4q21. The human genome contains numerous pairs of genes with similar “head-to-head” orientations and transcription start sites separated by less than 1 kb (Adachi and Lieber, 2002). Several of these gene pairs are known to be regulated by a single bidirectional promoter (Trinklein et al., 2004). Bidirectional promoters typically have high GC contents, frequently lack TATA boxes, and often are conserved among mouse orthologs (Orekhova and Rubtsov, 2013; Yang and Elnitski, 2008). These structural features are present in the bidirectional promoter region of THAP9/THAP9-AS1.

Bidirectional promoters regulate genes’ coordinated expression with complementary roles (Dovhey et al., 2000). They help maintain stoichiometric quantities of bidirectional gene pairs (Albig et al., 1997). They are also responsible for driving the transcription of genes involved in the same cellular pathway and regulating the expression of genes that need to be sequentially activated (Dovhey et al., 2000; Guarguaglini et al., 1997). In this regard, the bidirectional THAP9/THAP9-AS1 promoter region needs further experimental evidence to understand its regulatory nature in coordinating the expression THAP9 and THAP9-AS1.

It has been reported that THAP9 is a highly conserved gene, which has been identified in 178 organisms (Rashmi et al., 2021). The human THAP9 gene has 6 isoforms, out of which only one encodes for the full-length transposase protein. hTHAP9 belongs to the THAP (Thanatos-associated protein) protein family in humans, with twelve proteins (hTHAP0-hTHAP11) (Sabogal et al., 2010). Many THAP family proteins are known to be involved in human diseases. THAP1 has been associated with DYT6 dystonia (Sengel et al., 2011), THAP5 and THAP1 have been linked to apoptosis (Balakrishnan et al., 2009; Roussigne et al., 2003), LRRC49/THAP10 bidirectional gene pair are involved in breast cancer (De Souza Santos et al., 2008), and THAP11 has been implicated in the colon and gastric cancer (Parker et al., 2012; Zhang et al., 2020, p. 11). THAP9-AS1 (THAP9 antisense) is a newly annotated lncRNA coding gene by Ensembl that encodes 12 long non-coding RNAs. Meanwhile, recent reports have suggested that the THAP9-AS1 lncRNA has been implicated in pancreatic cancer, septic shock, and neutrophil apoptosis (Jia et al., 2019, p. 1; N. Li et al., 2020, p. 1). However, the roles of THAP9 and THAP9-AS1 in human pan-cancer are not well understood. This study identified the relationship between THAP9 and THAP9-AS1 expression and their possible roles in multiple cancer tumorigenesis models via a pan-cancer analysis of TCGA and GTEx databases.

Our gene expression analysis using TIMER2, EdgeR, and GEPIA2 suggests that abnormal expression of THAP9 and THAP9-AS1 frequently occurred in various types of cancer. We observed THAP9 be upregulated in CHOL, COAD, ESCA, LIHC, LUSC, LUAD, STAD, and THYM; downregulated in KIRC, KIRP, PRAD, THCA, TGCT, and UCEC. Moreover, THAP9-AS1 was upregulated in CHOL, THYM, DLBC, and PAAD, while it was downregulated in OV, SKCM, and THCA. We observed following the above results that THAP9 and THAP9-AS1 expressions were coordinately upregulated in CHOL and THYM compared with the corresponding normal tissues we used. However, THAP9 and THAP9-AS1 expression were coordinately downregulated in THCA. Therefore, independent and coordinated alteration in THAP9 and THAP9-AS1 expression in various cancers indicates that they may have different biological functions in different types of cancers. Regardless, aberrant expression levels of the THAP9 and THAP9-AS1 gene pair were associated with poor prognosis in many types of cancer, which strongly indicates their role as a potential prognostic biomarker in patients with cancer. Gene mutations also play an essential role in cancer pathogenesis. In this study, the most frequent DNA alterations of the THAP9 and THAP9-AS1 genes in the dataset from TCGA were amplifications.

Comprehensive gene expression studies provide an additional layer of information helpful in predicting gene function. The function of a gene can be implied by identifying genes that share its expression pattern. Genes that are turned on or turned off together under various conditions may encode proteins part of the same multiprotein machine or proteins involved in a complex coordinated activity. Characterizing an unknown gene’s function by grouping it with known genes that share its transcriptional behavior is sometimes called “guilt by association analysis (GBA)” (Alberts et al., 2002). GBA can be better explained by the concept that “a man is known by the company he keeps.” We can do GBA by performing gene coexpression analysis followed by constructing the gene interaction networks, often created by clustering associations from gene coexpression data. Genes that belong to the same cluster may be involved in common cellular pathways or processes. The study used the WGCNA method followed by Gene Ontology and KEGG pathway analysis using ShinyGO.

Interestingly, KEGG pathway analysis of genes coexpressed with THAP9 and THAP9-AS1 revealed the enrichment of pathways related to Herpes simplex virus 1 infection and several neurodegenerative disorders like Alzheimer’s, Parkinson’s, Huntington’s, etc. Several studies have reported Herpes simplex virus 1 infection as one of the causative agents of Alzheimer’s disease (Marcocci et al., 2020). Moreover, one study also reported the association of THAP9 in central nervous system tuberculosis. They observed a 5-fold upregulation of THAP9 in Tuberculous Meningitis (TBM) patients co-infected with HIV compared to patients with TBM (Kumar et al., 2012). This piques our speculation towards the possible role of THAP9 in Neurological Disorders. Enrichment of GO terms related to DNA binding, sequence-specific DNA binding, Transferase Complex, metal ion binding goes well with the fact that THAP9 is domesticated transposase and it consists of a characteristic THAP domain which is a zinc finger-type DNA binding domain that binds to a specific DNA sequence.

This study investigated the pan-cancer analysis of THAP9 and THAP9-AS1 gene pairs in various cancers. We explored the association of their aberrant expression with patient survival outcomes, followed by analyzing their functional association using gene coexpression and differential gene correlation analysis. This study has several limitations. First, although TCGA and GTEx datasets were included in this study, the numbers of each cancer type were still limited. Data about some particular cancer types were not available. Second, given the myriad individual differences among patients with cancer, it was challenging to cover all possible variations in this study. Finally, this study was only based on bioinformatics and relied on public databases. Future mechanistic studies to validate the expression and function of the two genes at the cellular and molecular levels are needed.

## Data Availability

Supplementary Data for the paper is available at https://doi.org/10.5281/zenodo.5867579.

## Abbreviations

ACC: adrenocortical carcinoma
BLCA: bladder urothelial carcinoma
BRCA: breast invasive carcinoma
CESC: cervical and endocervical cancers
CHOL: cholangiocarcinoma
COAD: colon adenocarcinoma
DLBC: lymphoid neoplasm diffuse large B-cell lymphoma
ESCA: esophageal carcinoma
GBM: glioblastoma multiforme
HNSC: head, and neck squamous cell carcinoma
KICH: kidney chromophobe
KIRC: kidney renal clear cell carcinoma
KIRP: kidney renal papillary cell carcinoma
LAML: acute myeloid leukemia
LGG: brain lower grade glioma
LIHC: liver hepatocellular carcinoma
LUAD: lung adenocarcinoma
LUSC: lung squamous cell carcinoma
MESO: mesothelioma
OV: ovarian serous cystadenocarcinoma
PAAD: pancreatic adenocarcinoma
PCPG: pheochromocytoma, and paraganglioma
PRAD: prostate adenocarcinoma
READ: rectum adenocarcinoma
SARC: sarcoma
SKCM: skin cutaneous melanoma
STAD: stomach adenocarcinoma
STES: stomach and esophageal carcinoma
TGCT: testicular germ cell tumors
THCA: thyroid carcinoma
THYM: thymoma
UCEC: uterine corpus endometrial carcinoma
UCS: uterine carcinosarcoma
UVM: uveal melanoma

## Materials and Methods

### 1. Analysis of promoter sequence

The promoter sequences were downloaded using EPDnew (Dreos et al., 2015), which is a new section under the well-known Eukaryotic Promoter Database (EPD) (https://epd.epfl.ch) (Dreos et al., 2015) that is an annotated non-redundant collection of eukaryotic POL II promoters where the transcription start site (TSS) has been determined experimentally. The core promoter elements in the region were identified using the Elements Navigation Tool (ElemeNT) (Sloutskin et al., 2015), a user-friendly web-based, interactive tool for predicting and displaying putative core promoter elements and their biologically-relevant combinations. ElemeNT’s predictions are based on biologically-functional core promoter elements and can infer core promoter compositions. The ElemeNT does not assume prior knowledge of the actual TSS position and can annotate any given sequence.

The bidirectional promoter region identified the CpG islands and other epigenetic marks using the UCSC Genome Browser (http://genome.ucsc.edu/). The CpG islands were plotted using EMBOSS Cpgplot (https://www.ebi.ac.uk/Tools/seqstats/emboss_cpgplot/).

### 2. Gene expression analysis

Differential gene expression analysis, to investigate the change in expression of specific genes in tumor vs. normal samples, was performed using TIMER2.0 (T. Li et al., 2020, p. 2), EdgeR (Robinson et al., 2010), and GEPIA2 tools (Tang et al., 2019, p. 2).

#### 2.1 TIMER2.0

TIMER2.0 (tumor immune estimation resource, version 2; http://timer.cistrome.org/) is a comprehensive resource for systematically analyzing differential gene expression between tumor and adjacent normal tissues. We used “THAP9” and “THAP9-AS1” as input in the “Gene_DE” module to evaluate the expression of the two genes in tumor tissue and adjacent normal tissues from 32 cancer types of the TCGA project (Tomczak et al., 2015). “THAP9-AS1” was not available on TIMER2.0.

#### 2.2 EdgeR

##### Sample Collection

We download HT-Seq-counts RNA-Seq data of 33 tumor types from the TCGA (The Cancer Genome Atlas) database (https://portal.gdc.cancer.gov/) using the GDC Data Transfer Tool (https://gdc.cancer.gov/access-data/gdc-data-transfer-tool). The dataset included the expression profiles of 60483 genes from 11,094 patients. Constraints used to generate the manifest file to be used with GDC Data Transfer Tool are as follows: Data Category: Transcriptome Profiling, Data Type: Gene-Expression quantification, Experimental strategy: RNA-Seq, Workflow type: HT-Seq-counts, Data format: txt, Access: open, Program: TCGA.

Details about the samples can be found in (**Figure 3b, Supplementary Table 3**). The row names of the downloaded HT-Seq-count matrix were Ensembl Gene Identifiers, and the column names represented the TCGA Sample IDs.

#### Sample Preprocessing

The Ensembl Gene Ids link to gene information in the Ensembl database (Howe et al., 2021). The “org.Hs.eg.db” R Bioconductor package (“org.Hs.eg.db,” n.d.) was used to convert the Ensembl Gene Ids to the Gene Symbols. Ensembl Ids that did not have an official gene symbol were dropped from the analysis. After filtering the samples, we were left with gene expression values of 34125 genes across 11,094 samples.

#### Differential Gene Expression Analysis

After preprocessing, we used the “edgeR” R Bioconductor package to store the data in a simple list-based data object called a DGEList for performing the differential gene expression analysis. This object is easy to use as it can be manipulated like an ordinary list in R, and it can also be subsetted like a matrix. The main components of the DGEList object were the matrix of read-counts and sample information in the “data.frame” format. Normalization by trimmed mean of M values (TMM) was performed with the “calcNormFactors” function in the “edgeR” package. The negative binomial (NB) distribution was used to model the RNA-seq read counts per gene per sample in edgeR. Then, estimating dispersion was calculated with the “estimateDisp” function. DEG analysis between each Tumor type (i.e., tumor versus normal) with more than 3 normal samples was performed using likelihood ratio tests (LRTs), including the “glmFit” and “glmLRT” functions in edgeR. Lastly, we plotted the fold change of THAP9 and THAP9-AS1 for comparison.

#### 2.3 GEPIA2

GEPIA2 (gene expression profiling interactive analysis, version 2; http://gepia2.cancer-pku.cn/#analysis) is an interactive web server for analyzing mRNA expression data from tumors and normal samples from TCGA (The Cancer Genome Atlas) and GTEx (Genotype-Tissue Expression) projects. We used the “Expression Analysis-Box Plots” module of GEPIA2 to obtain box plots of THAP9 and THAP9-AS1 expression between tumor and normal tissues. We set the P-value cutoff = 0.01, log2FC (fold change) cutoff = 1, and “Match TCGA normal and GTEx data”.

### 3. Prognostic analysis of THAP9 and THAP9-AS1

The GEPIA2 webserver (accessed 28 Nov 2021) was used to explore the prognostic values of THAP9 and THAP9-AS1 in different types of tumors in TCGA. The “Survival Map” module of GEPIA2 was used to obtain the overall survival (OS) and disease-free survival (DFS) significance map data with cutoff-high (50%) and cutoff-low (50%) values to split the high-expression and low-expression cohorts. The survival data was visualized with hazard ratio, 95% confidence intervals, and log-rank P values.

### 4. Mutation analysis in different types of tumors

The cBioPortal web server (http://www.cbioportal.org accessed 30 Nov 2021) (Cerami et al., 2012), is a comprehensive website, which explores, visualizes, and analyzes multidimensional cancer genomics data. We used the “Cancer Types Summary” module on the “TCGA PanCanAtlas” dataset available on the cBioPortal web server. Furthermore, the correlation between the genetic alteration of the two genes and their Overall survival prognosis was explored in the “Comparison” module. The prognostic values are presented with log-rank P values.

### 5. Guilt by Association Analysis

#### 5.1 Construction of Weighted Gene Co-Expression Network

GBA (guilt by association) analysis (Oliver, 2000) was used to identify co-expressing genes in each tumor and associated normal samples. The dataset used was the same as the one used for differential gene expression analysis using EdgeR (Section 2.2 Methods).

The gene expression values of each tumor and paired normal samples were subjected to the WGCNA, R Bioconductor package (Langfelder and Horvath, 2008) for weighted co-expression network construction. We used the “blockwiseModules” function in WGCNA, which performs automatic network construction and module detection on large expression datasets in a block-wise manner. In summary, it calculates the similarity matrix between each pair of genes across all samples based on its Pearson’s correlation value. Then, the similarity matrix is transformed into an adjacency matrix. Subsequently, the topological overlap matrix (TOM) and the corresponding dissimilarity (1-TOM) value are computed. Finally, a dynamic tree cut (DTC) algorithm detects gene co-expression modules. The signed modules were constructed with a cut height of 0.995 and a minimum module size of 30 genes. Then we used the modules associated with THAP9 and THAP9-AS1 in Cytoscape (Shannon et al., 2003) to visualize the top 20 genes co-expressed with THAP9 and THAP9-AS1 in each tumor vs. normal pair.

#### 5.2 Gene ontology (GO) and KEGG pathway enrichment analysis

GO analysis (Ashburner et al., 2000) is a helpful method for annotating genes and gene sets with biological characteristics for high-throughput genome or transcriptome data. The Kyoto Encyclopedia of Genes and Genomes (KEGG) pathway (Kanehisa and Goto, 2000) is a knowledge base for systematic analysis of gene functions. GO, and KEGG pathway enrichment analysis was performed for genes part of the THAP9 and THAP9-AS1 co-expression module using the “ShinyGO” webserver (Ge et al., 2020). P-value cutoff (FDR) of 0.05 was set as the cut-off criterion for extracting the top 10 enriched GO terms (Biological Process (BP), Cellular Component (CC), and Molecular Function (MF)) and KEGG pathways. Further, we merged all the GO-BP, GO-CC, GO-MF, and KEGG pathways enriched in all the tumor and normal samples and used the “word cloud” package in python to visualize the overall enrichment GO and KEGG pathways in Normal vs. tumor samples.

#### 5.3 Differential Correlation analysis

For the differential co-expression analysis between the tumor and normal samples in each cancer, R-package DGCA version 1.0.2 (McKenzie et al., 2016) was used. Firstly, we checked the correlation between THAP9 and THAP9-AS1 in each tumor vs. normal pair using “plotCors” function in DGCA, which uses “Pearson” correlation by default. Following that, genes that are differentially correlated with THAP9 and THAP9-AS1 were computed with the “ddCorAll” function using “corrType” as spearman correlation. This pipeline provided the Spearman coefficient and the corresponding p values for each pair of genes across samples. Significant changes in differential correlation between the two conditions (tumor vs. normal) were then identified using a Fisher’s Z-test. The correlation between THAP9/THAP9-AS1 and other genes was classified as having a gain of correlation or loss of correlation, and based upon the threshold for correlation significance; the gene pairs were grouped into nine different correlation classes (+/+; +/−; +/0; −/+; −/0; −/−; 0/+; 0/0; 0/−). The classes show correlation as positive (+), negative (−), or not significant (0) for each gene and condition when contrasting the groups (tumor/normal). GO term enrichment analysis of differential correlation-classified genes was performed using the DGCA function “ddcorGO.”.that

